# Meaning Guides Attention in Real-World Scene Images: Evidence from Eye Movements and Meaning Maps

**DOI:** 10.1101/207076

**Authors:** John M. Henderson, Taylor R. Hayes

**Author notes:** Correspondence: John M. Henderson, Center for Mind and Brain, 267 Cousteau Place, University of California, Davis, CA 95618.

## Abstract

We compared the influences of meaning and salience on attentional guidance in scene images. Meaning was captured by “meaning maps” representing the spatial distribution of semantic information in scenes. Meaning maps were coded in a format that could be directly compared to maps of image salience generated from image features. We investigated the degree to which meaning versus image salience predicted human viewers’ spatial distribution of attention over scenes, with attention operationalized as duration-weighted fixation density. The results showed that both meaning and salience predicted the distribution of attention, but that when the correlation between meaning and salience was statistically controlled, meaning accounted for unique variance in attention but salience did not. This pattern was observed for early as well as late fixations, for fixations following short as well as long saccades, and for fixations including or excluding the centers of the scenes. The results strongly suggest that meaning guides attention in real world scenes. We discuss the results from the perspective of a cognitive relevance theory of attentional guidance in scenes.

## Introduction

We can only attend to a fraction of the visual stimulation available to us at any given moment. For this reason, visual attention is guided through scenes in real time, with the eyes shifting position about three times each second on average to select informative objects and scene regions for scrutiny (Buswell, 1935; Hayhoe & Ballard, 2005; Henderson, 2003, 2017; Henderson & Hollingworth, 1999; Land & Hayhoe, 2001; Rayner, 2009; Yarbus, 1967). How does the brain determine which scene regions and elements should be attended at any given moment?

Most recent research on attentional guidance in real world scene images has focused on the idea that attention is primarily driven by low-level image features. Image guidance theory has its roots in models of attention and visual search that focus on the attraction of attention by primitive visual features and feature differences (Treisman & Gelade, 1980; Wolfe & Horowitz, 2017). When applied to real world scenes, the most influential instantiation of this type of theory is based on image salience, which proposes that saliency maps are generated by pooling contrasts in semantically uninterpreted image features such as luminance, color, and edge orientation (Borji, Parks, & Itti, 2014; Borji, Sihite, & Itti, 2013; Harel, Koch, & Perona, 2006; Itti & Koch, 2001; Parkhurst, Law, & Niebur, 2002). In this theoretical approach, regions that are uniform along these features are considered uninformative, whereas those that differ from neighboring regions across spatial scales are taken to be worthy of attention. That is, salient points in the map serve as a prediction about the spatial distribution of attention in a scene, and these points often correlate with observed human fixations. In this view, attentional guidance is fundamentally a reaction to image features in the scene, with attention captured or “pulled” to visually salient scene regions (Henderson, 2007). An appeal of image guidance theory based on salience is that image salience is both neurobiologically inspired and computationally tractable (Henderson, 2017). The saliency map approach has served an important heuristic function in the study of attention and eye movements in scene perception by providing an explicit model that generates quantitative predictions about attention and eye movements.

Despite the substantial influence of image saliency on research in scene perception, it is also well established that semantic content of a scene and the viewers task also influence viewing (Buswell, 1936; Yarbus, 1967). Indeed, when directly tested, image salience often does a poor job of accounting for attention in real-world scene viewing (Einhäuser, Rutishauser, & Koch, 2008; Henderson, Brockmole, Castelhano, & Mack, 2007; Henderson, Malcolm, & Schandl, 2009; Tatler, Hayhoe, Land, & Ballard, 2011; Underwood, Foulsham, & Humphrey, 2009). To account for these observations, cognitive guidance models place primary emphasis on cogitive control of attention. In this view, attention is “pushed” by the cognitive system to scene regions that are semantically informative and cognitively relevant in the current situation (Henderson, 2007). For example, in the cognitive relevance model (Henderson et al., 2007, 2009), attention is guided by semantic representations that code the meaning of the scene and its local regions (objects, surfaces, and other interpretable entities) with respect to the viewer’s current goals and task (Buswell, 1935; Hayhoe & Ballard, 2005; Hayhoe, Shrivastava, Mruczek, & Pelz, 2003; Henderson, 2003, 2007, 2017; Henderson & Hollingworth, 1999; Land & Hayhoe, 2001; Rothkopf, Ballard, & Hayhoe, 2007; Tatler et al., 2011; Turano, Geruschat, & Baker, 2003; Võ & Wolfe, 2013; Yarbus, 1967). The cognitive relevance model posits that the representations used to assign task relevance and meaning for attentional priority encode knowledge about the world itself (world knowledge), as well as knowledge about the general scene concept (scene schema knowledge) and the current scene instance (episodic scene knowledge) (Henderson & Ferreira, 2004; Henderson & Hollingworth, 1999).

Most proponents of image guidance acknowledge that meaning must play some role in attentional guidance. Nevertheless, much of the research on attentional guidance in real-world scene images has been motivated by and focused on image salience as instantiated by saliency maps. One reason for this emphasis is the relative tractability of image salience; it is far easier to quantifying image features than it is to quantify meaning (Henderson, 2017). To investigate meaning and to compare its influence to salience, it is necessary to represent both constructs so that comparable quantitative predictions can be generated from them.

To provide a method for directly comparing the influences of meaning and salience on the guidance of attention, we have recently developed the concept of *meaning maps* (Henderson & Hayes, 2017). Meaning maps draw inspiration from two classic scene viewing studies (Antes, 1974; Mackworth & Morandi, 1967). In these studies, images were divided into regions and subjects were asked to rate each region based on how easy that region would be to recognize (Antes, 1974) or how informative it was (Mackworth & Morandi, 1967). In both studies, the eye movements of a different group of subjects were measured while they viewed the rated images. In general, viewers looked more at the more highly rated regions. We modified and extended these methods to develop meaning maps for real world scenes. We used crowd-sourced responses in which we asked naïve subjects to rate the meaningfulness of a large number of scene patches. Specifically, photographs of scenes were divided into a dense array of objectively defined circular overlapping patches at two spatial scales (Figure 1). These patches were then presented to raters independently of the scenes from which they are taken and raters were asked to indicate how informative or recognizable they judged it to be (Figure 2). Finally, we constructed smoothed maps for each scene based on interpolated ratings over a large number of raters (Figure 3). The basic idea of the meaning map is that it captures the spatial distribution of the semantic content of a scene in the same format as a saliency map captures the spatial distribution of image salience. Like image salience, meaning is spatially distributed non-uniformly across scenes, with some scene regions relatively rich in semantic content and others relatively sparse.

**Figure 1.**
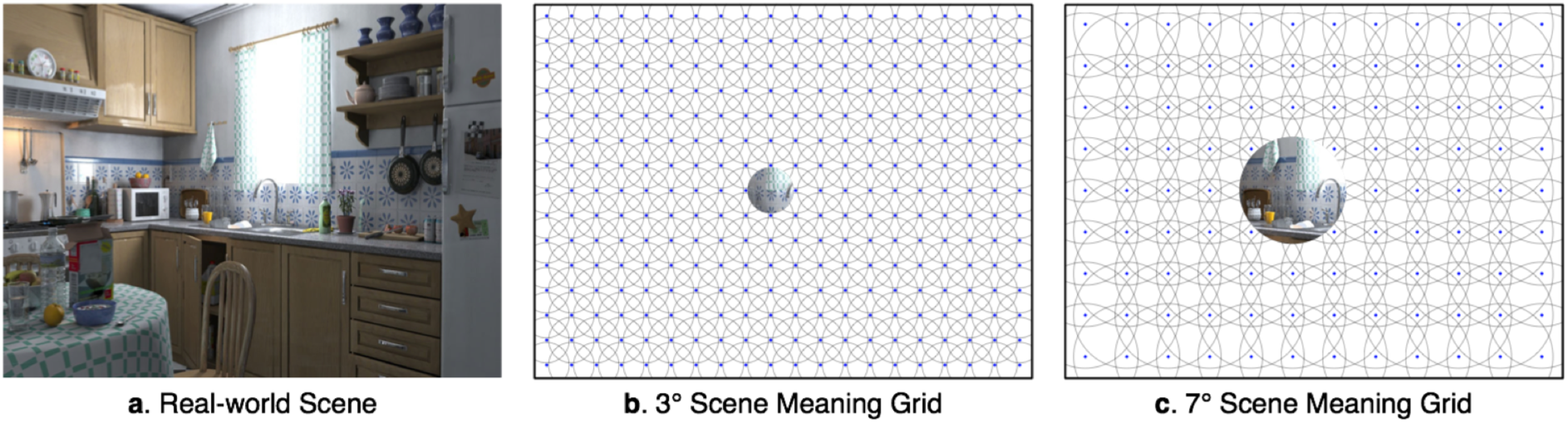
Real-world scene and corresponding tiled patch grids. (a) Example real-world scene. (b) Overlapping circular patches used for meaning rating at 3° and (c) at 7° spatial scales. The blue dots in (b) and (c) denote the center of each circular patch and the image circles show examples of the content captured by the 3◦ and 7◦ scales for the example scene

**Figure 2.**
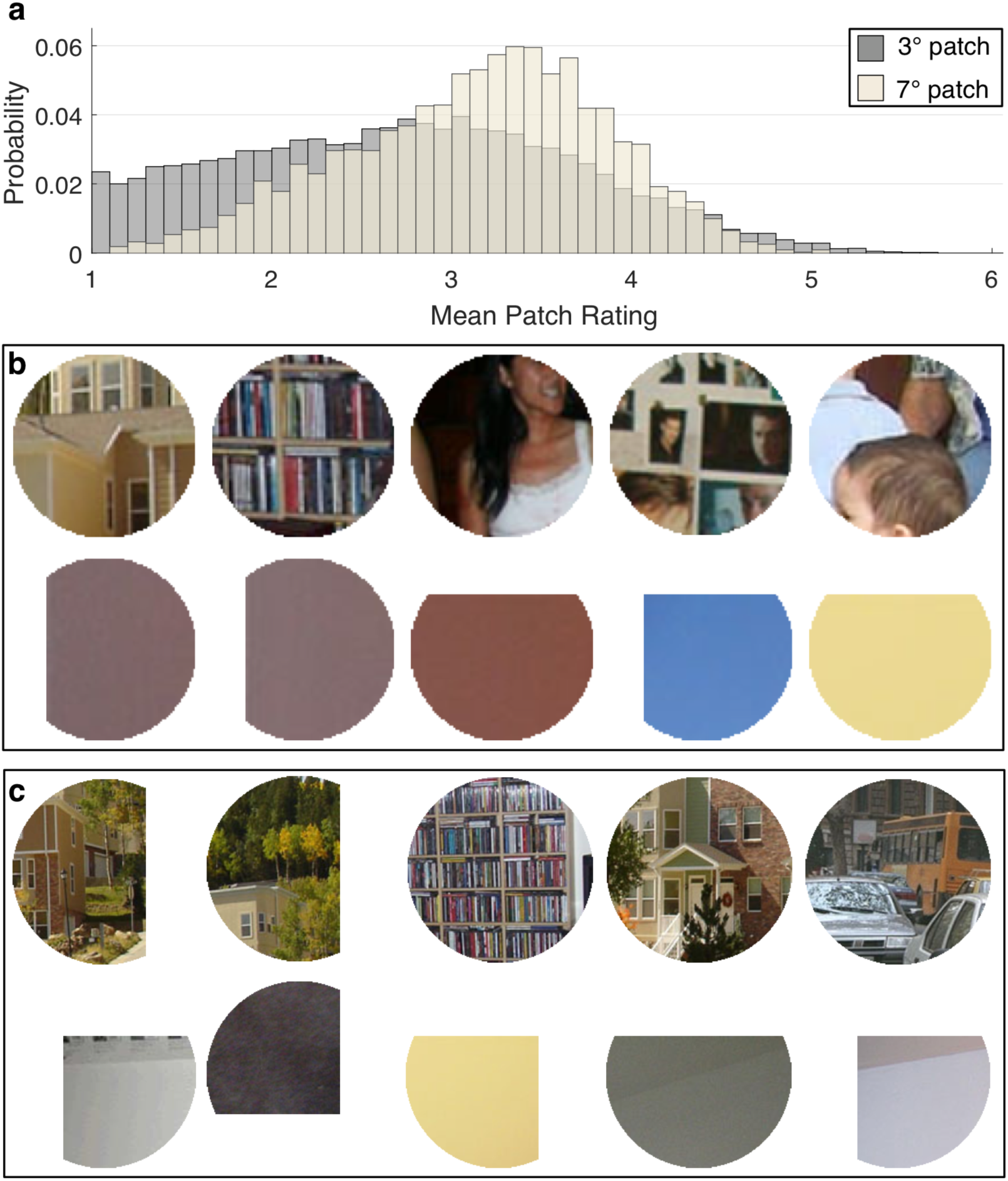
Rating distributions and example high and low patches. (a) Distribution of ratings for 3 and 7 patches across all raters and scenes. (b) Example highest and lowest rated nonoverlapping patches for 3° and (c) 7° patches.

**Figure 3.**
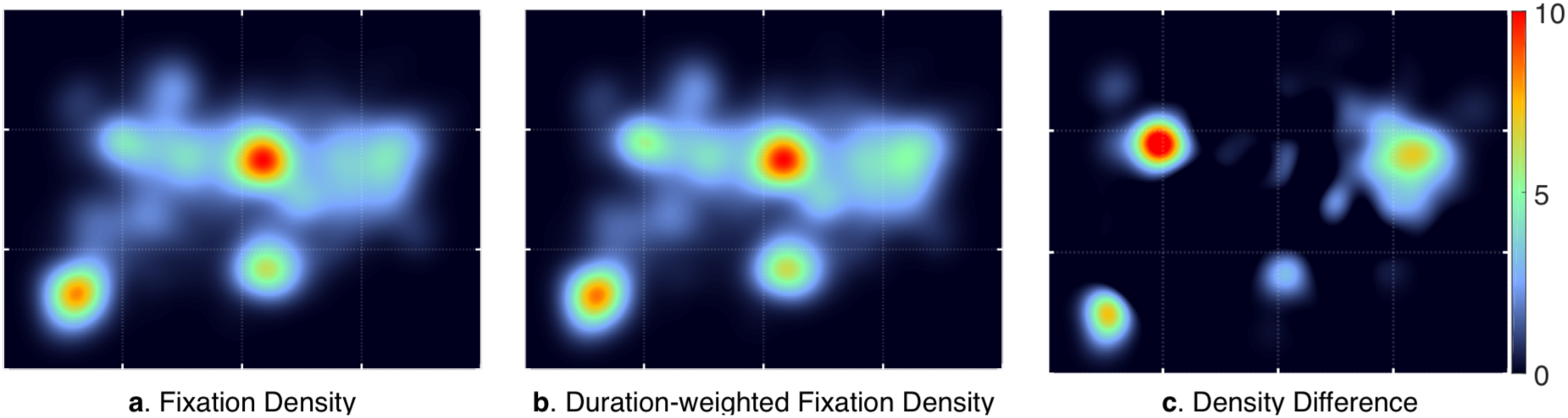
Duration-weighted fixation density. Example (a) fixation density and (b) durationweighted fixation density, for all fixations on one scene. (c) The density difference depicting the absolute-value difference in the two densities, with hotter regions representing greater difference.

Because meaning maps are represented in the same format as saliency maps, they can be directly compared to saliency maps. A meaning map provides the conceptual analog of a saliency map by capturing the spatial distribution of semantic features (rather than image features) across a scene. They can be used to generate predictions concerning attentional guidance using the same methods that have been used to test the goodness of fit of predictions from saliency theory (Carmi & Itti, 2006; Itti, Koch, & Niebur, 1998; Parkhurst et al., 2002; Torralba, Oliva, Castelhano, & Henderson, 2006). And the predictions for attentional guidance generated from meaning maps can be compared to those generated from saliency maps. In short, meaning maps and saliency maps provide a foundation for directly contrasting the influences of meaning and salience on attentional guidance.

In an initial study, we investigated the relative ability of meaning maps and saliency maps to predict attentional guidance during scene viewing (Henderson & Hayes, 2017). In that study and in keeping with the literature on scene perception, attention maps were based on the locations of eye fixations. We found that both meaning and salience could predict the distribution of attention over scenes, with meaning accounting for more variance in attention than image salience. However, we also found that meaning and salience were themselves highly correlated. Furthermore, when the variance due to salience was controlled, meaning accounted for a significant amount of the remaining variance in attention, but when meaning was controlled, no further variance in attention was accounted for by salience. These data held for both early and later fixations during viewing, including the very earliest fixations on the scenes. The data strongly suggested that attention is guided by meaning rather than saliency.

The present study was designed to extend the original meaning map results. A potential concern with the original report is that the attention maps were based on fixation locations that did not take into account fixation durations (Henderson & Hayes, 2017). The fixation location analysis was an important first step because the research assessing saliency maps to date has similarly focused on fixation location (Borji et al., 2014, 2013; Harel et al., 2006; Itti & Koch, 2001; Parkhurst et al., 2002). However, fixation durations vary, and this variability reflects a variety of factors including attention related to perceptual and cognitive processing. When more attention is needed on an object or other scene entity, fixations are directed to that entity for more time (Einhäuser & Nuthmann, 2016; Henderson, Nuthmann, & Luke, 2013; Henderson, Weeks, & Hollingworth, 1999; Henderson & Pierce, 2008; Henderson & Smith, 2009; Laubrock, Cajar, & Engbert, 2013; Nuthmann, 2017; Nuthmann, Smith, Engbert, & Henderson, 2010). The distribution of attention over a scene therefore depends on both the location and duration of attentional selection (Henderson, 2003). For this reason, we report here a new set of analyses designed to determine how well meaning and salience predict attentional guidance in scenes account for how long attention is focused at each location.

In summary, the goal of this study was to test current theoretical approaches to attentional guidance in real-world scenes. We applied our recently developed method, meaning maps, to capture the spatial distribution of semantic content across scenes. We then tested cognitive and image guidance theories by comparing the ability of meaning maps and saliency maps to predict attentional guidance during real world scene viewing, with attention operationalized as the duration-weighted fixations of subjects viewing the scenes.

## Method

### Meaning Maps

For this study we used the meaning maps developed in Henderson and Hayes (2017).

#### Subjects

Scene patch ratings were performed by 165 subjects on Amazon Mechanical Turk. Subjects were recruited from the United States, had a hit approval rate of 99% and 500 hits approved, and were only allowed to participate in the study once. Subjects were paid $0.50 cents per assignment and all subjects provided informed consent.

#### Stimuli

Stimuli were 40 digitized (1024x768 pixels) photographs of real world scenes depicting a variety of indoor and outdoor environments. The full set of scene images can be found in the supplementary materials of Henderson and Hayes (2017). Each scene was decomposed into a series of partially overlapping (tiled) circular patches at spatial scales of 3° and 7° (Figure 1). Simulated recovery of known scene properties (e.g., luminance) indicated that the underlying property could be recovered well (98% variance explained) using these patches (see Appendix), suggesting that this method is sufficiently sensitive to underlying scene structure. The full patch stimulus set consisted of 12000 unique 3° patches and 4320 unique 7° patches for a total of 16320 scene patches.

#### Procedure

Each subject rated 300 random patches extracted from 40 scenes. Subjects were instructed to assess the meaningfulness of each patch based on how informative or recognizable they thought it was. Subjects were first given examples of two low-meaning and two high-meaning scene patches to make sure they understood the rating task, and then rated the meaningfulness of scene patches on a 6-point Likert scale (’very low’, ’low’, ’somewhat low’, ’somewhat high’, ’high’, ’very high’). Patches were presented in random order and without scene context, so ratings were based on context-free judgments. Each unique patch was rated 3 times by 3 independent raters for a total of 48960 ratings. However, due to the high degree of overlap across patches, each patch contained rating information from 27 independent raters for each 3° patch and 63 independent raters for each 7° patch. Figure 2 shows the distribution of ratings and the highest and lowest rated non-overlapping patches across all scenes at the two patch sizes. The lowest rated patches tended to come from the edges of the pictures, which accounts for their truncated shapes.

Meaning maps were generated from the ratings by averaging, smoothing, and then combining 3° and 7° maps from the corresponding patch ratings. The ratings for each pixel at each scale in each scene were averaged, producing an average 3° and 7° rating map for each scene. The average 3° and 7° rating maps were then smoothed using thin-plate spline interpolation (Matlab ’fit’ using the ’thinplateinterp’ method). Finally, the smoothed 3° and 7° maps were combined using a simple average, i.e., (3° map + 7° map)/2). This procedure was used to create a meaning map for each scene. The final map was blurred using a Gaussian kernel followed by a multiplicative center bias operation which down-weighted the scores in the periphery to account for the central fixation bias, the commonly observed phenomenon in which subjects concentrate their fixations more centrally and rarely fixate the outside border of a scene (Borji et al., 2013; Henderson et al., 2007; Tatler, 2007). This center bias operation is also commonly applied to saliency maps.

To investigate the relationship between the generated meaning maps and image-based saliency maps, saliency maps for each scene were computed using the Graph-based Visual Saliency (GBVS) toolbox with default settings (Harel et al., 2006). GBVS is a prominent saliency model that combines maps of neurobiologically inspired low-level image features. The same center bias operation described for the meaning maps was applied to the saliency maps to down-weight the periphery.

#### Histogram Matching

Meaning and saliency maps were normalized to a common scale using image histogram matching, with the duration-weighted fixation map for each scene serving as the reference image for the corresponding meaning and saliency maps. Histogram matching of the meaning and saliency maps was accomplished using the Matlab function ’imhistmatch’ in the Image Processing Toolbox.

### Eyetracking Experiment and Attention Maps

#### Subjects

Seventy-nine University of South Carolina undergraduate students with normal or corrected-to-normal vision participated in the experiment. All subjects were naive concerning the purposes of the experiment and provided informed consent. The eye movement data from each subject was inspected for excessive artifacts caused by blinks or loss of calibration due to incidental movement by examining the mean percent of signal across all trials using Matlab. Fourteen subjects with less than 75% signal were removed, leaving 65 subjects for analysis who tracked very well (mean signal = 91.74%). We have previously used this corpus to investigate individual differences in scan patterns in scene perception (Hayes & Henderson, 2017) as well as for the initial study of meaning maps (Henderson & Hayes, 2017).

#### Apparatus

Eye movements were recorded with an EyeLink 1000+ tower mount eyetracker (spatial resolution 0.01) sampling at 1000 Hz (SR Research, 2010b).

Subjects sat 90 cm away from a 21” monitor, so that scenes subtended approximately 33°x25° of visual angle at 1024x768 pixels. Head movements were minimized using a chin and forehead rest. Although viewing was binocular, eye movements were recorded from the right eye. The experiment was controlled with SR Research Experiment Builder software (SR Research, 2010a).

#### Stimuli

Stimuli consisted of the 40 digitized photographs of real-world scenes that were used to create the meaning and saliency maps.

#### Procedure

Subjects were instructed to view each scene in preparation for a later memory test. The memory test was not administered. Each trial began with fixation on a cross at the center of the display for 300ms. Following central fixation, each scene was presented for 12s while eye movements were recorded. Scenes were presented in the same order for all subjects.

A 13-point calibration procedure was performed at the start of each session to map eye position to screen coordinates. Successful calibration required an average error of less than 0.49° and a maximum error of less than 0.99°. Fixations and saccades were segmented with EyeLink’s standard algorithm using velocity and acceleration thresholds (30/s and 9500°/s; SR Research, 2010b).

Eye movement data were imported offline into Matlab using the EDFConverter tool. The first fixation, always located at the center of the display as a result of the pretrial fixation period, was eliminated from analysis.

#### Attention maps

The distribution of attention over a scene is a function of the locations and durations of eye fixations (Henderson, 2003). Although maps created from fixation locations alone (Henderson & Hayes, 2017) and from the duration-weighted fixations were similar, they were not identical (see also Henderson, 2003). An example of the difference can be seen in Figure 3 by comparing fixation density maps based on location alone (Figure 3a) to maps of location weighted by duration (Figure 3b). The difference in the two maps is shown in Figure 3c, with regions of greater difference shown with hotter colors. As can be seen, some regions changed their relative attentional weighting when duration was considered. For the present analyses, we therefore created attention maps from fixation density weighted by fixation duration.

To create duration-weighted attention maps, a duration weight was generated for every fixation following the initial (experimenter defined) fixation. Because average fixation durations vary reliably and systematically across subjects (Castelhano & Henderson, 2008a; Henderson & Luke, 2014; Rayner, Li, Williams, Cave, & Well, 2007), duration weights were based on subject-normalized values. We first generated each subject’s fixation duration distribution across all 40 scenes. We then defined 2 parameters for these distributions, an upper bound 95th percentile cutoff (any values in the 95 percentile received a weight value of 1.0) and the lower bound minimum weight cutoff of 0.1 (any value below the 0.1 percentile received a weight value of 0.1 to avoid weights of 0). Each fixation was therefore weighted from 0.1 to 1.0 based on its place in the overall distribution. Fixation-weighted values were accumulated across all subjects adding the weight to each location, producing a weighted fixation frequency matrix for each scene. Finally, a Gaussian low pass filter with a circular boundary and a cutoff frequency of -6dB was applied to the matrix for each scene to account for foveal accuity/eye tracker error. The Gaussian low-pass function is from the MIT Saliency Benchmark code (https://github.com/cvzoya/saliency/blob/master/code_forMetrics/antonioGaussian.m). With a cut off frequency (fc=6) the window size is approximately 2 degrees of visual angle. An example of a resulting duration-weighted attention map is shown in Figure 4c.

**Figure 4.**
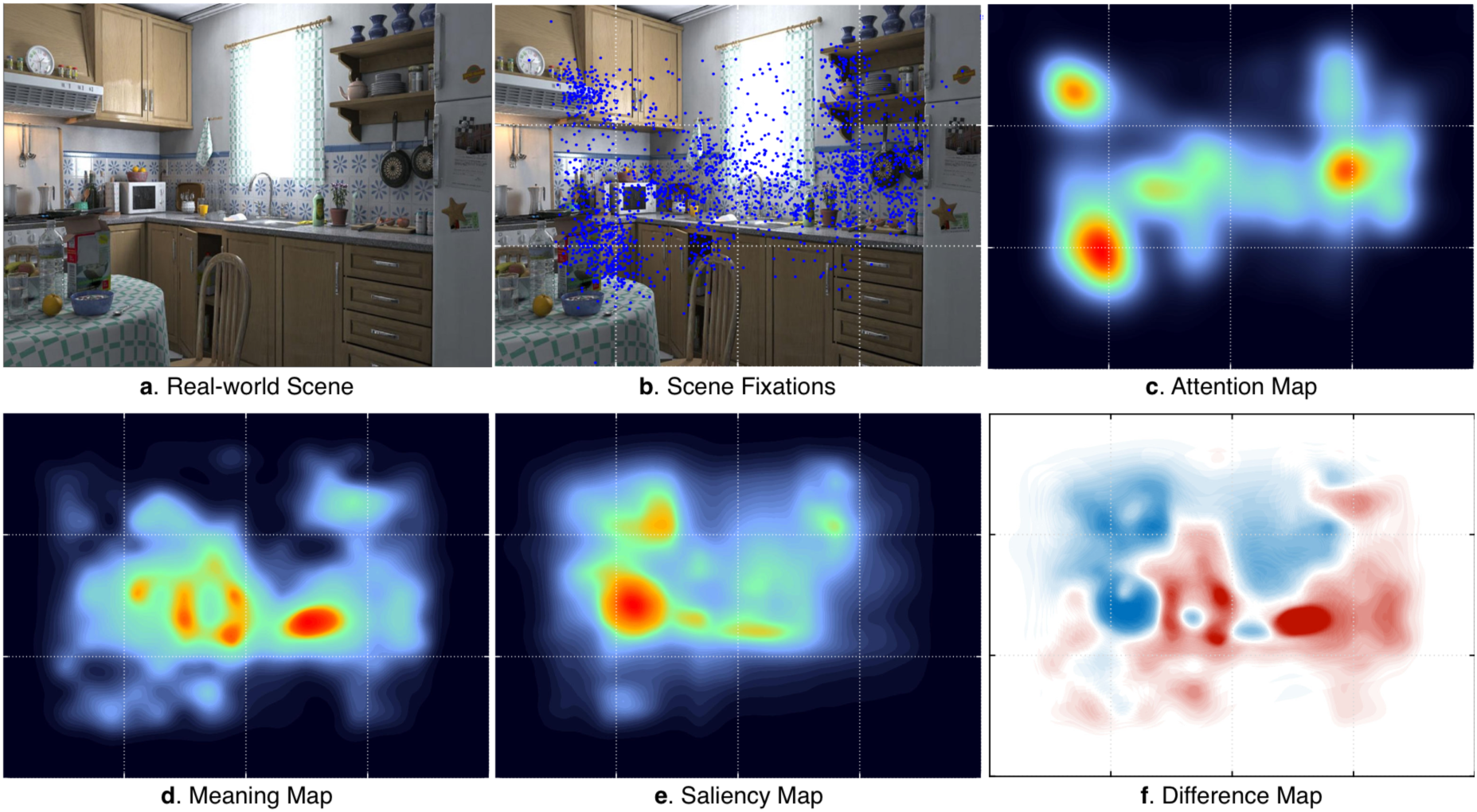
Example data used in the analyses. (a) Real-world scene, (b) viewers’ fixations superimposed on the scene as blue dots, (c) the duration-weighted attention map derived from the fixations, (d) meaning map, (e) saliency map for the example scene, and (f) the difference between the meaning and saliency maps, with regions of greater meaning shown in red and greater saliency shown in blue. The meaning and saliency maps are on the same scale (0-1) and the difference map is on a 10% scale (0-0.10). Note that the guidelines in the scene were not shown to subjects and are presented to facilitate comparison across panels.

## Results

We can take meaning maps and saliency maps as predictions concerning how viewers will distribute their attention over scenes. To investigate how well meaning maps and saliency maps predict the distribution of attention, it is important to assess the degree of association between the meaning maps and saliency maps themselves. For the scenes used here, the correlation between meaning and salience was 0.80 averaged across the 40 scenes (Henderson & Hayes, 2017). This correlation is consistent with the suggestion that attention effects that have previously been attributed to salience could be due to meaning (Henderson et al., 2007, 2009; Nuthmann & Henderson, 2010). At the same time, meaning and salience did not share 36% of their variance, and we can ask how well this unshared variance in each predicts attention.

The critical empirical question in the present study was how well the two types of prediction maps (meaning and saliency maps) capture the distribution of attention. To investigate this question, we used linear correlation (Bylinskii, Judd, Oliva, Torralba, & Durand, 2016) to determine the degree to which meaning maps (Figure 4d) and saliency maps (Figure 4e) statistically predicted the spatial distribution of attention (Figure 4b) as captured by the duration-weighted attention maps (Figure 4c). This method allows us to assess the degree to which meaning maps and saliency maps account for shared and unique variance in the attention maps. There are many ways in which prediction maps can be tested against attention maps, and no method is perfect (Bylinskii et al., 2016). We chose here a map-level analysis method that makes relatively few assumptions, is intuitive, can be visualized, generally balances the various positives and negatives of different analysis approaches, and that allows us to tease apart variance due to salience and meaning.

Figure 5 presents the primary data for each of the 40 scenes. Each data point shows the relationship (R^2^ value) between the meaning map and the observed attention map for each scene (red), and between the saliency map and the observed attention map for each scene (blue). The top half of Figure 5 shows the squared linear correlations. On average across the 40 scenes, meaning accounted for 50% of the variance in fixation density (M=0.50, SD=0.12) and salience accounted for 35% of the variance in fixation density (M=0.35, SD=0.12). A two-tailed t-test revealed this difference was statistically significant, *t*(78) = 5.38, *p* < .0001, 95% CI [0.09,0.20].

**Figure 5.**
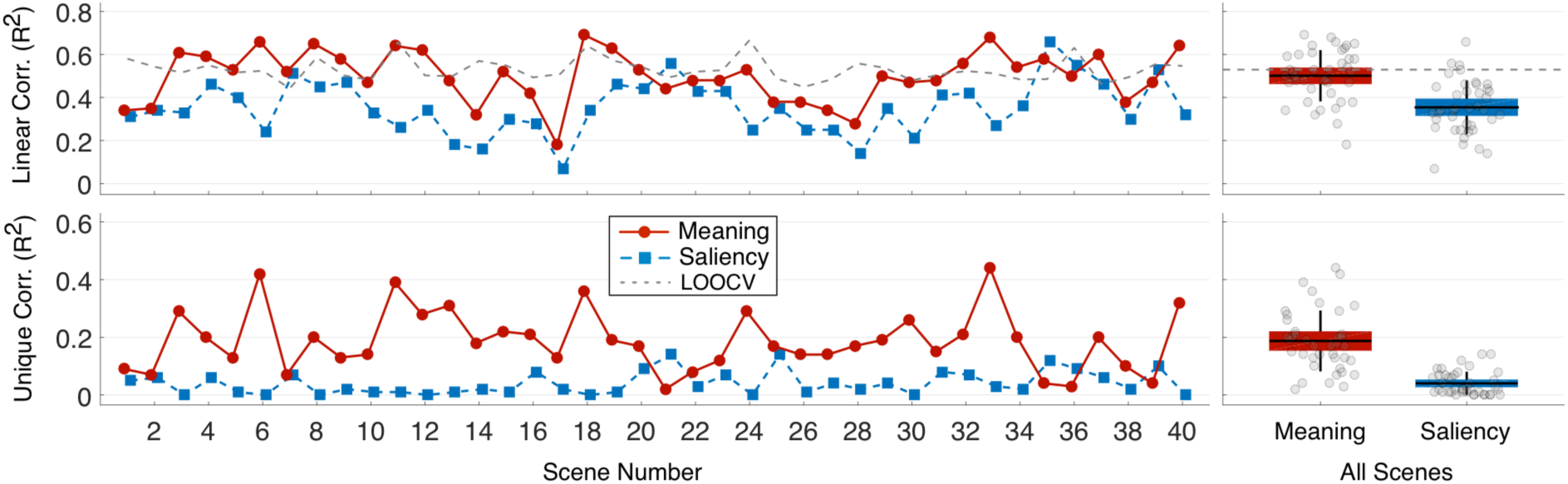
Squared linear correlation and semi-partial correlation by scene and across all scenes. The line plots show the linear correlation (top) and semi-partial correlation (bottom) between durationweighted fixation density and meaning and salience by scene. The scatter box plots on the right show the corresponding grand mean (black horizontal line), 95% confidence intervals (colored box), and 1 standard deviation (black vertical line) for meaning and salience across all 40 scenes. The dotted lines in the top panels show consistency across subjects based on leave-one-out cross-validation (LOOCV).

We used leave-one-out cross-validation to estimate the upper limit on salience and meaning performance given subject variability in attention (Torralba et al., 2006). The cross-validation analysis sets an expected maximum in the ability of meaning and salience to account for attention. Specifically, we computed for each scene a duration-weighted group attention map as described above for 64 subjects and a test map for the 65th subject. The linear correlation of the group and test maps was computed, and this was repeated for all 65 subjects. Mean correlations by scene and across scenes were then generated. The results are shown as dotted lines in the top panels of Figure 5, with the left panel showing the mean linear correlation for each scene and the right panel showing the grand mean across scenes. Across all scenes, cross-validation R^2^ was 0.53 (SD=0.05). Meaning map performance was not statistically different from this theoretical maximum, as revealed by a two-tailed t-test, t(78) = 1.37, p=0.17, 95% CI [-0.01,0.07]. In comparison, saliency maps produced poorer performance than the theoretical maximum, t(78) = 8.17, p < .0001, 95% CI [0.13,0.22]. These results show that meaning maps accounted for attention about as well as possible given the reliability of the subject data, whereas saliency maps performed significantly below this level.

To examine the unique variance in attention explained by meaning and salience when controlling for their shared variance, we computed squared semi-partial correlations (bottom half of Figure 5). Across the 40 scenes, meaning accounted for a significant 19% additional variance in the attention maps after controlling for salience (M=0.19, SD=0.11), whereas salience accounted for a non-significant 4% additional variance after controlling for meaning (M=0.04, SD=0.04). A two-tailed t-test confirmed that this difference was statistically significant, *t*(78) = 8.22, *p* < .0001, 95% CI [0.11, 0.18]. These results show that meaning explained the distribution of attention over scenes better than salience. It has sometimes been proposed that during scene viewing, attention is initially guided by image salience, but that as viewing progresses over time, meaning begins to play a greater role (Henderson & Ferreira, 2004; Henderson & Hollingworth, 1999; Mannan, Ruddock, & Wooding, 1996; Parkhurst et al., 2002). To test this proposal, we conducted temporal time-step analyses. Linear correlation and semi-partial correlations were conducted based on a series of attention maps, with each map generated from each sequential eye fixation (i.e., 1^st^, 2^nd^, 3^rd^ fixation, etc.) in each scene. This method allowed us to test whether the relative importance of meaning and salience in predicting attention changed over time. The results are shown in Figure 6. For the linear correlations, the relationship was stronger between the meaning and attention maps for all time steps (top of Figure 6) and was highly consistent across the 40 scenes. Meaning accounted for 33.0%, 32.1%, and 29.7% of the variance in the first 3 fixations, whereas salience accounted for only 9.5%, 15.2%, and 16.6% of the variance in the first 3 fixations, respectively. Two sample two-tailed t-tests were performed for all 38 time points, and p-values were corrected for multiple comparisons using the false discovery rate (FDR) correction (Benjamini & Hochberg, 1995). This procedure confirmed the advantage for meaning over salience at all 38 time points (*FDR* < 0.05).

**Figure 6.**
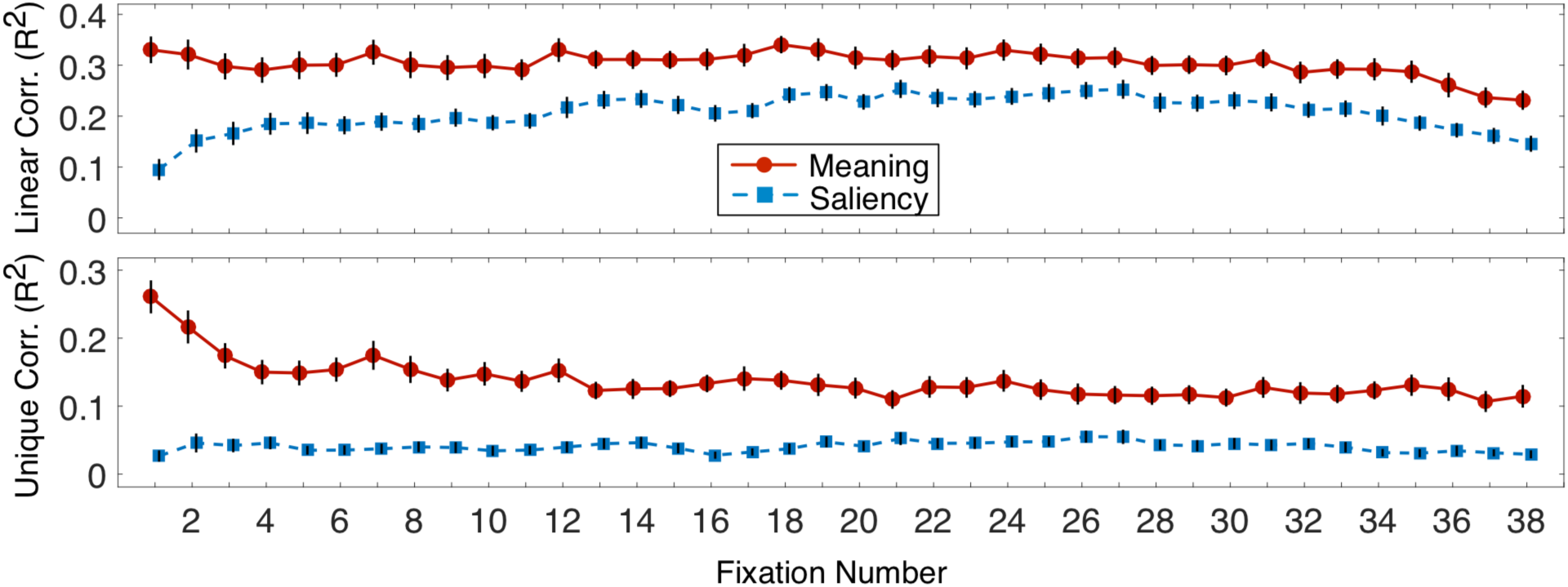
Squared linear correlation and squared semi-partial correlation as a function of fixation number. The top panel shows the squared linear correlation between duration-weighted fixation density and meaning and salience as a function of fixation order across all 40 scenes. The bottom panel shows the corresponding semi-partial correlation as a function of fixation order across all 40 scenes. Error bars represent standard error of the mean.

The improvement in R^2^ for the meaning maps over saliency maps observed in the overall analyses was again found to hold across all 38 time steps in the partial correlations (bottom of Figure 6) (*FDR* < 0.05), with meaning accounting for 26.1%, 21.7%, and 17.4% of the unique variance in the first 3 fixations, whereas salience accounted for 2.7%, 4.6%, and 4.2% of the unique variance in the first 3 fixations, respectively. In conclusion, counter to the salience-first hypothesis but consistent with results based on unweighted fixations reported (Henderson & Hayes, 2017), in both the correlation and semi-partial correlation analyses meaning accounted for more variance in attention than salience from the very first fixation. These results indicate that meaning begins guiding attention as soon as a scene appears (Rider, Coutrot, Pellicano, Dakin, & Mareschal, 2018).

### Central Region Knock-Out Analyses

It is commonly found in eyetracking studies that viewers tend to concentrate their fixations near the center of a real world scene and rarely fixate the outside borders of a scene (Borji et al., 2013; Henderson et al., 2007; Tatler, 2007). As noted in the methods, in creating the final meaning maps, we used a multiplicative center bias operation to down-weight the scores in the periphery and consequently up-weight the center, as is commonly done with saliency maps. However, to further ensure that the advantage of meaning maps over saliency maps in predicting the distribution of attention was not due to a center bias advantage for the meaning maps, we also conducted additional analyses in which the data from the central 7° of each map (attention, meaning, and saliency) were removed. Differences in the success of meaning and saliency maps in this analysis therefore can not be due to differences in the ability of meaning maps to predict central fixations since they are no longer included. The results of these analyses were qualitatively and quantitatively very similar to the complete analyses.

Figure 7 presents the linear correlation data used to assess the degree to which meaning maps and saliency maps accounted for shared and unique variance in the attention maps for each scene excluding the central 7°. Each data point shows the R^2^ value for the prediction maps (meaning and saliency) and the observed attention maps for saliency (blue) and meaning (red). The top of Figure 7 shows the squared linear correlations. On average across the 40 scenes, meaning accounted for 46% of the variance in fixation density (M=0.46, SD=0.11) and saliency account for 34% of the variance in fixation density (M=0.34, SD=0.13). A two-tailed t-test revealed this difference was statistically significant, *t*(78) = 4.39, *p* < .0001, 95% CI [0.06, 0.17].

**Figure 7.**
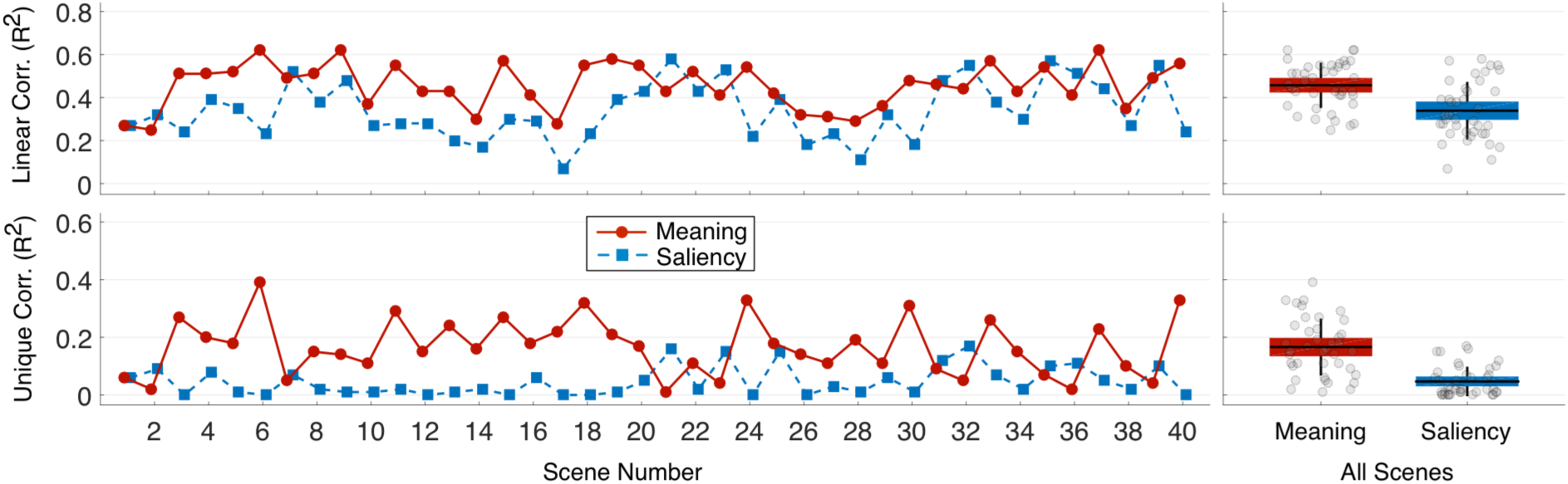
Squared linear correlation and semi-partial correlation by scene and across all scenes with 7° *center removed.* The line plots show the linear correlation (top) and semi-partial correlation (bottom) between duration-weighted fixation density and meaning and salience by scene after removing the central 7° from each scene. The scatter box plots on the right show the corresponding grand mean (black horizontal line), 95% confidence intervals (colored box), and 1 standard deviation (black vertical line) for meaning and salience across all 40 scenes.

To examine the unique variance in attention explained by meaning and salience excluding the central 7° and when controlling for their shared variance, we computed squared semi-partial correlations. These correlations, shown in the bottom of Figure 7, revealed that across the 40 scenes, meaning captured more than 3 times as much unique variance (M=0.17, SD=0.10) as saliency (M=0.05, SD=0.05). A two-tailed t-test confirmed that this difference was statistically significant, *t*(78) = 6.78, *p* < .0001, 95% CI [0.08, 0.16]. These results confirm those of the complete analysis and indicate that meaning was better able than salience to explain the distribution of attention over scenes even when the central 7° of maps was removed.

To test whether the overall advantage of meaning over salience early in viewing was due to meaning at the center, we conducted the fixation series analysis excluding the central 7° of maps. Figure 8 shows the temporal time-step analyses with the central 7° of maps removed. Linear correlation and semi-partial correlation were conducted as in the main time-step analyses based on a series of attention maps generated from each sequential eye fixation in each scene. Using the same testing and false discovery rate correction as in the main analyses, 34 of 38 time points were significantly different in both the linear and semi-partial analyses (*FDR* < 0.05), excluding fixations 21, 25, 27, and 28. Importantly for assessing initial control of attention during scene viewing, in the linear correlation analysis (top of Figure 8), meaning accounted for 22.9%, 27.0%, and 26.7% of the variance in the first three fixations, whereas salience accounted for only 10.2%, 14.9%, and 16.2% of the variance in the first three fixations. Critically, when controlling for the correlation among the two prediction maps with semi-partial correlations, the advantage for the meaning maps observed in the overall analyses was also found to hold across all time steps, as shown in the bottom of Supplementary Figure 8 (*FDR* < 0.05). Meaning accounting for 17.9%, 17.8%, and 15.7% of the unique variance in the first 3 fixations, whereas salience accounted for 5.2%, 5.6%, and 5.4% of the unique variance in the first three fixations, respectively. Consistent with the overall correlation and semi-partial correlation analyses, meaning produced an advantage over salience from the very first fixation even when the central region of each map was removed from the analysis. These results indicate that when overt attention leaves the center of a scene, meaning guides even those earliest shifts of overt attention. These results are especially strong evidence for the control of attention by meaning because removing the central 7° should disadvantage the meaning maps because photographers tend to center meaningful information in photographs (Tatler, 2007). Nevertheless, the meaning maps continued to outperform the saliency maps in both the overall variance and unique variance accounted for in the attention maps.

**Figure 8.**
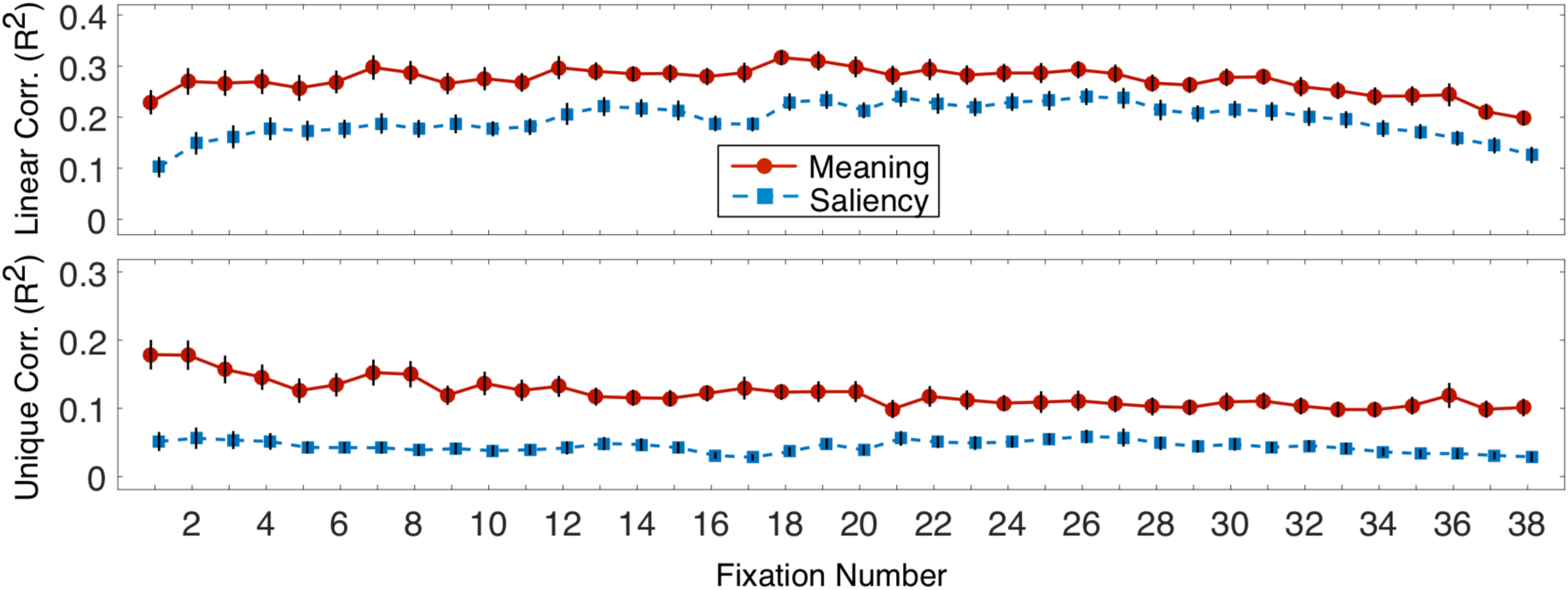
Squared linear correlation and squared semi-partial correlation as a function of fixation number with 7° center removed. The top panel shows the squared linear correlation between fixation density and meaning (red) and salience (blue) as a function of fixation order averaged over all 40 scenes. The bottom panel shows the corresponding semi-partial correlation as a function of fixation order averaged over all 40 scenes. Error bars represent standard error of the mean.

### Saccade Amplitude Analyses

It could be that meaning controls attention as it is guided within objects and nearby scene regions, but that salience controls attention as it is guided from one scene region to another. If this is true, then meaning should be more highly related to attentional selection following shorter saccades, whereas image salience should be more highly related of attention following longer saccades. To investigate this prediction, we conducted an analysis in which we examined how meaning and salience related to attention following saccades of different amplitudes.

Figure 9 presents the distribution of saccade amplitudes in the present study. The average amplitude was 3.5 degrees, but as typically observed in scene viewing, saccade amplitude varied considerably (Henderson & Hollingworth, 1999). Once again, we used correlation analyses to assess the degree to which meaning maps and saliency maps accounted for shared and unique variance in the attention maps data for fixations following saccades of different amplitudes. For these analyses saccade amplitudes were binned by percentile. Each data point shows the R^2^ value for the observed attention maps for saliency (blue) and meaning (red) at each saccade amplitude decile. The middle of Figure 9 shows the squared linear correlations and the bottom of Figure 9 shows the unique variance accounted for by meaning and salience. The R^2^ values for meaning and salience differed for all amplitudes except the very longest decile in both figures (*FDR* < .05). These results confirm those of the complete analysis and indicate that meaning was better able than salience to explain the distribution of attention over scenes even when attention was not limited to the object or scene region at the current point of attention.

**Figure 9.**
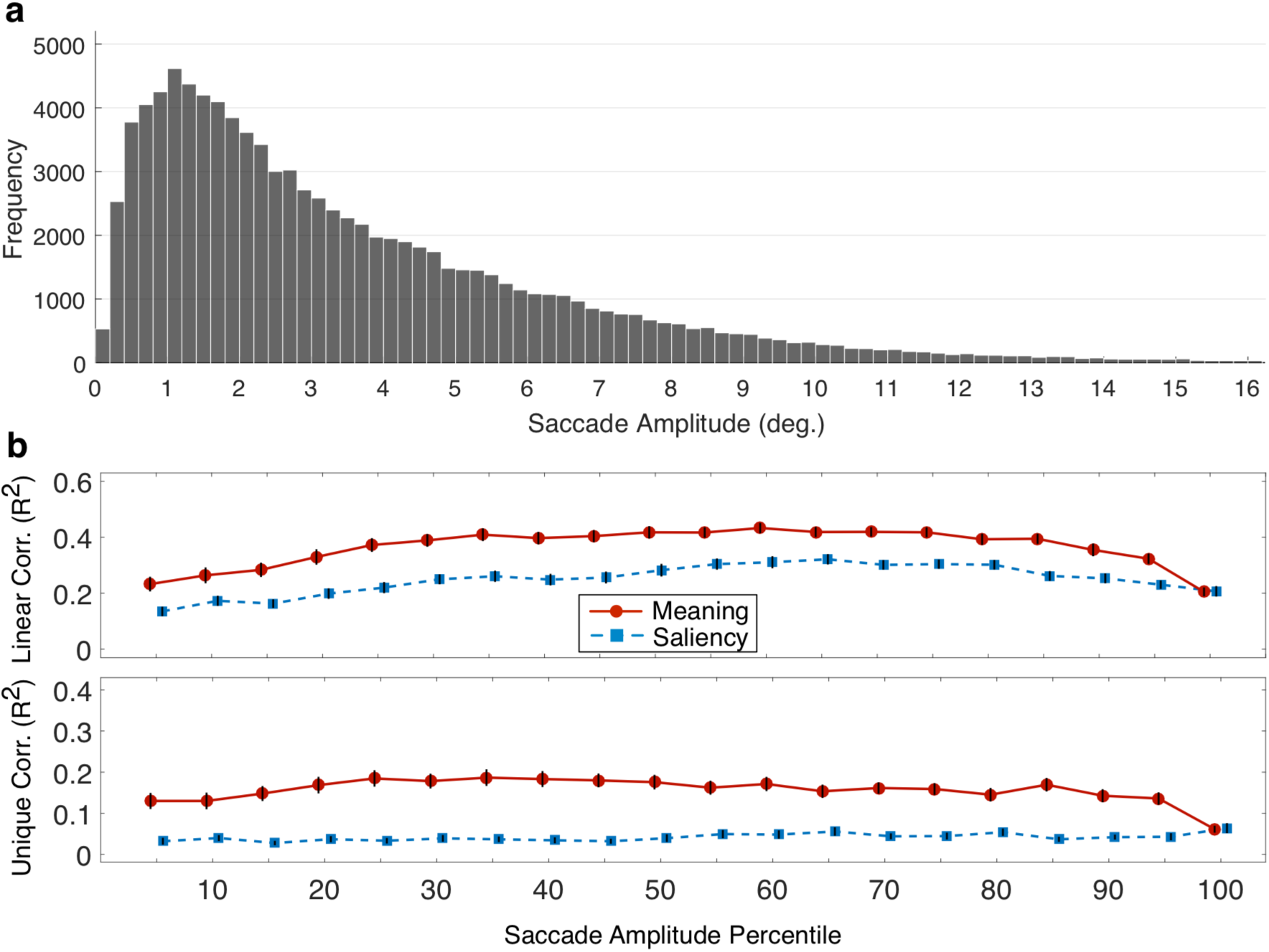
Squared linear correlation and squared semi-partial correlation as a function of saccade amplitude to fixation. (a) The distribution of saccade amplitudes observed in the experiment. (b) The squared linear correlations between duration-weighted fixation density for meaning and salience as a function of the saccade amplitude percentiles prior to fixation. (c) The corresponding semi-partial correlations as a function of saccade amplitude. Data points are averages across all 40 scenes. Error bars represent standard error of the mean.

## General Discussion

Image salience as instantiated by computationally derived saliency maps currently provides a central theoretical framework and empirical paradigm for understanding how attention is guided through real-world scenes. Yet human viewers are known to be highly sensitive to the semantic content of the visual world that they perceive, suggesting that attention may be directed by semantic content rather than image salience. Until recently it has been difficult to directly contrast the influence of image salience and meaning. To address this difficulty, we developed a new method for identifying and representing the spatial distribution of meaning in any scene (Henderson & Hayes, 2017). The resulting “meaning maps” quantify the spatial distribution of semantic content across scenes in the same format that saliency maps quantify the spatial distribution of image salience. Meaning maps therefore provide a method for disentangling the distribution of meaning from the distribution of image salience. In the present study, we used meaning maps to test the relative importance of meaning and salience during scene viewing by testing meaning maps and saliency maps against observed duration-weighted attention maps.

The results showed that both meaning maps and saliency maps were able to account for considerable variance in attention maps, suggesting that they both offered good predictions concerning attention. However, meaning maps and saliency maps are themselves strongly correlated (Henderson & Hayes, 2017). When these correlations were statistically controlled, meaning maps accounted for additional unique variance in the duration-weighted distribution of attention over scenes. On the other hand, the variance due to visual salience was completely accounted for by meaning, such that saliency maps accounted for no additional unique variance in the attention maps when the variance accounted for by meaning was controlled. These results suggest that meaning plays the primary role in directing attention through scenes.

A similar dominance of meaning over salience was observed throughout the viewing period, with unique variance accounted for by meaning beginning with the first subject-determined fixation. Contrary to salience-first models, these results suggest that meaning influences attentional guidance more strongly than salience both early and later during scene viewing. The results indicate that meaning begins guiding attention as soon as a scene appears, and suggest that viewers are able to determine very quickly (within the first glimpse) where meaningful regions within the current scene are to be found and to direct their attention based on that assessment.

The strong role of meaning in guiding attention in scenes can be accommodated by a theoretical perspective that places explanatory primacy on scene semantics. For example, on the *cognitive relevance* model (Henderson et al., 2007, 2009), the priority of an object or scene region for attention is determined solely by its meaning in the context of the scene and the current goals of the viewer, and not by its visual features or salience. On this model, meaning determines attentional priority, with image properties used only to generate perceptual objects and other perceptually based potential saccade targets. Critically, then, attentional priority is assigned to potential attentional targets not based image saliency, but rather based on knowledge representations (e.g., knowledge about what objects are likely to be present and where those objects are likely to be found). In this model, the visual stimulus is relevant in that it is used to generate perceptual objects and other targets for attention, and processes related to salience may be relevant in determining whether a perceptual object is generated, but the image features themselves provide a flat (that is, unranked) landscape of potential attentional targets rather than a landscape ranked by salience (Henderson et al., 2007). Instead, on this model, knowledge representations provide the attentional priority ranking to the targets based on their meaning (Henderson, 2003; Henderson et al., 2007, 2009).

It is important to note that the cognitive relevance model does not require meaning be assigned simultaneously across the entire scene to all perceptually mapped potential saccade targets. That is, the model does not require a strong “late-selection” view of scene perception in which all objects and scene regions are fully identified before they are attended. There are two reasons for this. First, when a scene is initially encountered, the “gist” of the scene can be quickly apprehended (Biederman, 1972; Castelhano & Henderson, 2008b; Fei-Fei, Iyer, Koch, & Perona, 2007; Potter, 1975) and can guide attention at the very earliest points of scene viewing (Castelhano & Henderson, 2003; Henderson & Hollingworth, 1999; Oliva & Torralba, 2006; Võ & Henderson, 2010). Apprehending the gist allows access to schema representations that provide constraints on what objects are likely to be present and where those objects are likely to be located (Henderson, 2003; Henderson & Hollingworth, 1999). Information retrieved from memory schemas can be combined with low-quality peripheral visual information from the periphery to assign tentative meaning to perceptual objects and other scene regions. These initial representations provide a rich set of priors and can be used to generate predictions for guiding attention to regions that have not yet been identified (Henderson, 2017). Second, as shown in the present study as well as many others (Henderson et al., 1999), most saccades during scene viewing are relatively short, with the average amplitude of about 3.5° in the present study. The implication is that attention is frequently guided from the current location to the next location based on information that is relatively close to the fovea, where identity and meaning can easily be ascertained. Extraction of meaning from nearby regions cannot be the entire story for attentional guidance given that meaning continues to dominate salience even for fixations following longer saccades, as shown in the present study, but it does suggest that for the many shorter shifts of attention, meaning is at least partly derived from a spatially local semantic analysis of the scene. For longer saccades, it is likely that guidance is based on scene representations retrieved from memory as described above.

The present results at first glance appear to be at odds with past studies that have shown correlations between visual salience and attention. How can we account for this discrepancy? One explanation can be found in the strong correlation between meaning and visual salience. We have hypothesized in the past that this correlation is likely to be high (Henderson et al., 2007). Meaning maps provide a method for testing hypothesis, and robust support was found for it, with an average correlation of 0.80 between meaning and salience (Henderson & Hayes, 2017). Given this correlation, salience can do a reasonably good job of predicting meaning-driven attention. From an engineering perspective, this might be sufficient. However, from the perspective of the neuro-cognitive study of human vision in which the goal is to provide a theoretical account of how the brain guides attention, a focus on salience will be misleading. Instead, the present results along with previous results (Henderson & Hayes, 2017) strongly suggest that meaning, not visual salience, is the causal factor that guides attention.

### Limitations and Future Directions

We note several limitations and caveats of this study and our earlier meaning map investigation (Henderson & Hayes, 2017). First, we have so far used a single viewing task. It has been shown that attention as indexed by eye movements differs over the same scene depending on the task (Castelhano, Mack, & Henderson, 2009; Henderson et al., 1999; Mills, Hollingworth, Van der Stigchel, Hoffman, & Dodd, 2011; Yarbus, 1967), and it could be that under other task instructions, image salience would play a greater role than meaning. While this is a possibility, the memorization task used here is a relatively unstructured free-viewing task in which viewers are not explicitly or implicitly directed to meaningful scene regions. Therefore, this task would not seem to favor meaning-based over image-based attentional guidance. Nevertheless, we can not rule out the possibility that salience might play a more important role in other tasks, and it will be important to assess the relative influences of meaning and salience in guiding attention in different viewing tasks.

Second, although meaning was the stronger predictor of attention on average and for the majority of scenes (36 out of 40 scenes) tested here, salience did perform better for four scenes. The question arises why these four scenes showed the opposite pattern. One possibility is that there may simply be statistical noise in one or more of the maps (meaning, saliency, or attention) for a given scene that occasionally leads to a random reversal of the true pattern. Another possibility is that there is some systematic difference in the scenes that show the reversed pattern. We were not able to discern any particular regularities across those scenes, but a future direction for study will be to compare different classes of scenes (e.g., indoor versus outdoor; natural versus man-made) to determine whether meaning and salience play greater or lesser roles for specific types of scenes.

Third, in the present study we defined meaning in a context-free manner, in the sense that each scene patch was rated for meaning without regard to the scene it came from. Meaning could instead be defined in a context-dependent manner, with the meaning of a scene region assessed in terms of its scene context. Similarly, meaning could vary as a function of the viewer’s task. So far we have focused on context-free meaning as a first step, but it will be important to determine how meaning changes as the context changes, and in turn how context-dependent meaning influences attention. One way to determine context-dependent meaning is to ask participants to indicate directly (e.g., via mouse click) which regions in a scene they find most interesting (Onat, Açık, Schumann, & König, 2014). However, subjects might click on regions that their attention has been drawn to, potentially confounding visual salience and meaning and leading to some circularity in using these clicks to predict future attention. Alternatively, consistent with the present approach, we might ask subjects to rate independent experimenter-defined scene patches but within the context of the entire scene or a specific task.

Fourth, we have chosen here to compare meaning to the class of saliency models that are inspired and motivated by neurobiologically plausible assumptions about the nature of visual computation in the human visual system (Borji & Itti, 2013; Itti & Koch, 2001; Itti et al., 1998). This class of saliency model continues to inspire a vast amount of research across many disciplines. Within this class of model, we have used the graph-based visual saliency (GBVS) implementation because it is typically the best performer (Walther & Koch, 2006). Indeed, in our own comparisons of saliency models, GBVS outperforms other similar models on our dataset. However, it should be noted that another class of model based on learning within deep neural networks (DNNs) has recently been advanced as a competitor to traditional saliency models (Vig, Dorr, & Cox, 2014). For example, DeepGaze II, the current top performer in this class, learns where people attend in scenes from training sets of fixations over object features and then predicts fixations on new scenes (Kummerer, Wallis, Gatys, & Bethge, 2017). Interesting issues for future research include comparison of predictions from current DNNs and meaning maps, and extending DNNs to include meaning. However, an important consideration from the perspective of understanding human neuro-cognitive processes is whether these models trade neurobiological plausibility and transparency for engineering expediency.

## Conclusion

In this study we employed recently developed methods for comparing the relationship between the spatial distribution of meaning and image salience to the spatial distribution of attention in scene viewing (Henderson & Hayes, 2017). We investigated the relative importance of meaning and salience on the guidance of attention in scenes as indexed by attention maps based on duration-weighted fixations. We found that the spatial distribution of meaning was better able than image salience to account for the guidance of attention, both overall and when controlling for the correlation of meaning and salience. Furthermore, we found that the advantage of meaning over image salience appeared from the very beginning of scene viewing, held over both shorter and longer shifts of attention, and persisted when the central region of each scene was removed from analysis. This pattern of results is consistent with a cognitive relevance theory of scene viewing in which attentional priority is assigned to scene regions based on semantic information rather than visual salience.

## Acknowledgements

We thank the members of the UC Davis Visual Cognition Research Group for their feedback and comments. Research reported in this publication was supported by the National Science Foundation under award number BCS-1636586, and by the National Eye Institute of the National Institutes of Health under award number R01EY027792. The content is solely the responsibility of the authors and does not necessarily represent the official views of the National Institutes of Health.

## Appendix

### Patch Density Parameter Estimation

The optimal meaning map grid density for each patch size was estimated by simulating the recovery of known image properties (i.e., luminance and entropy). For the sake of simplicity and visualization, the simulation procedure will be described in terms of luminance recovery, but the same procedure was also applied to edge density and entropy recovery.

The first step in the recovery simulation was to generate the ground truth luminance image for each scene for a given patch size, which sets an upper limit on the luminance resolution that can be recovered. The ground truth luminance image for each scene was computed by taking the scene luminance image and convolving it with a circular mean mask for a given patch size (i.e., 3° and 7°). Then, the patch density grid (simulating patch ratings) was systematically varied from 50 to 1000 patches (3°) and 40 to 200 (7°) and recovery of the ground truth was performed for each grid. The recovery procedure consisted of taking the mean of each patch from the original luminance image and then using thin plate interpolation to interpolate between the patches across each grid. If the patch density was low enough that the entire image was not tiled, then the background was set to the mean value across all the patch samples in the grid.

Figure A1 shows an example of the recovery procedure for the scene shown in Figure 1a for a patch density of 88 (a) and 300 (b). As can be seen by comparing the ground truth (left) to the interpolated recovery (right), a patch density of 300 provides an excellent estimate of the ground truth. Figure A2 shows luminance, edge density, and entropy recovery (R^2^) for the 3° patch size (a) and the 7° patch size (b) as a function of patch density. Recovery improvement plateaus at a patch density of 300 patches for the 3° patch size and 108 patches for the 7° patch size.

**Figure A1.**
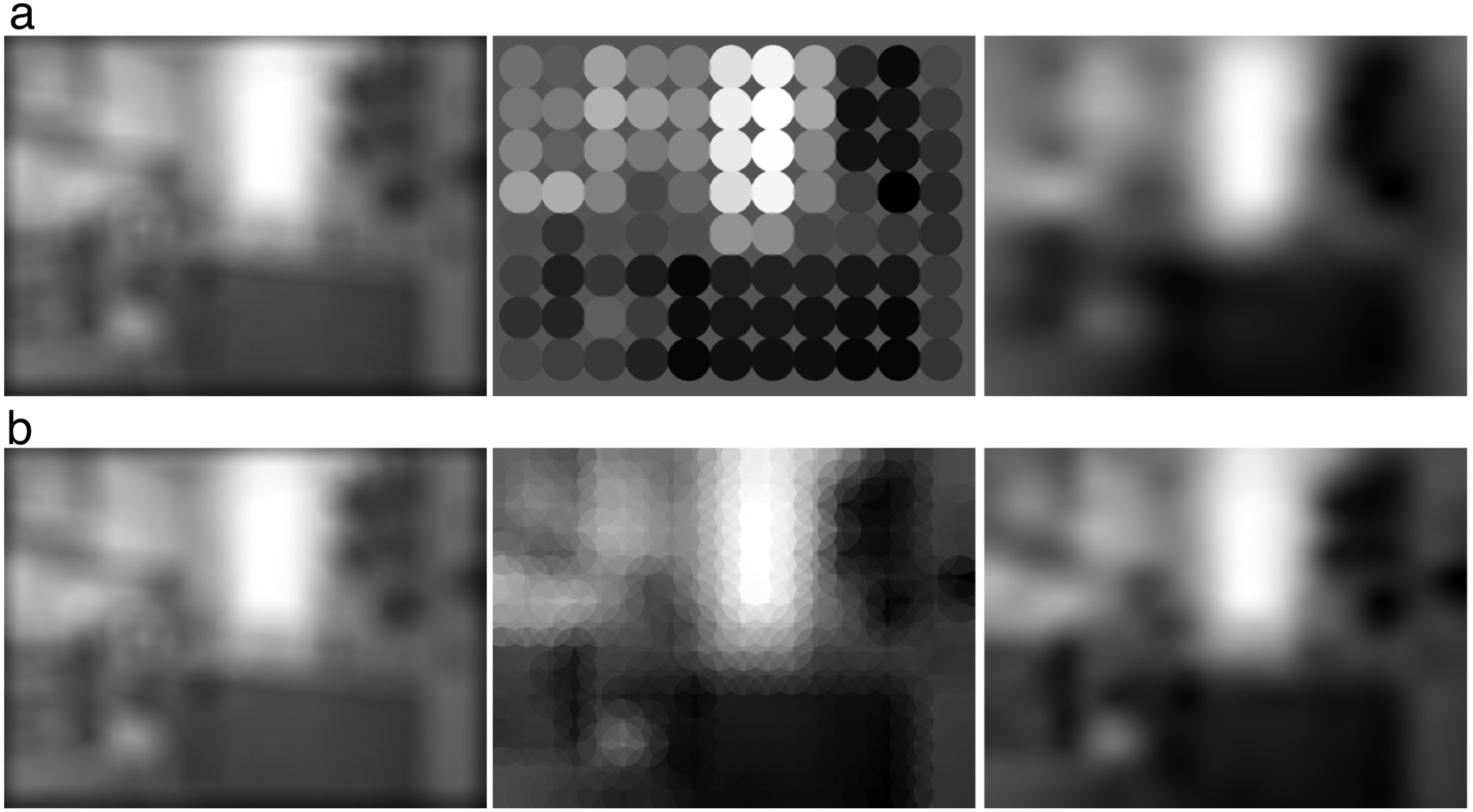
Example of scene luminance recovery. From left to right, the 3° ground truth luminance, simulated rating density, and interpolated recovery images are shown for a patch density of 88 (a) and a patch density of 300 (b). A comparison of the ground truth and recovery indicates that a patch density value of 300 provided excellent recovery.

**Figure A2.**
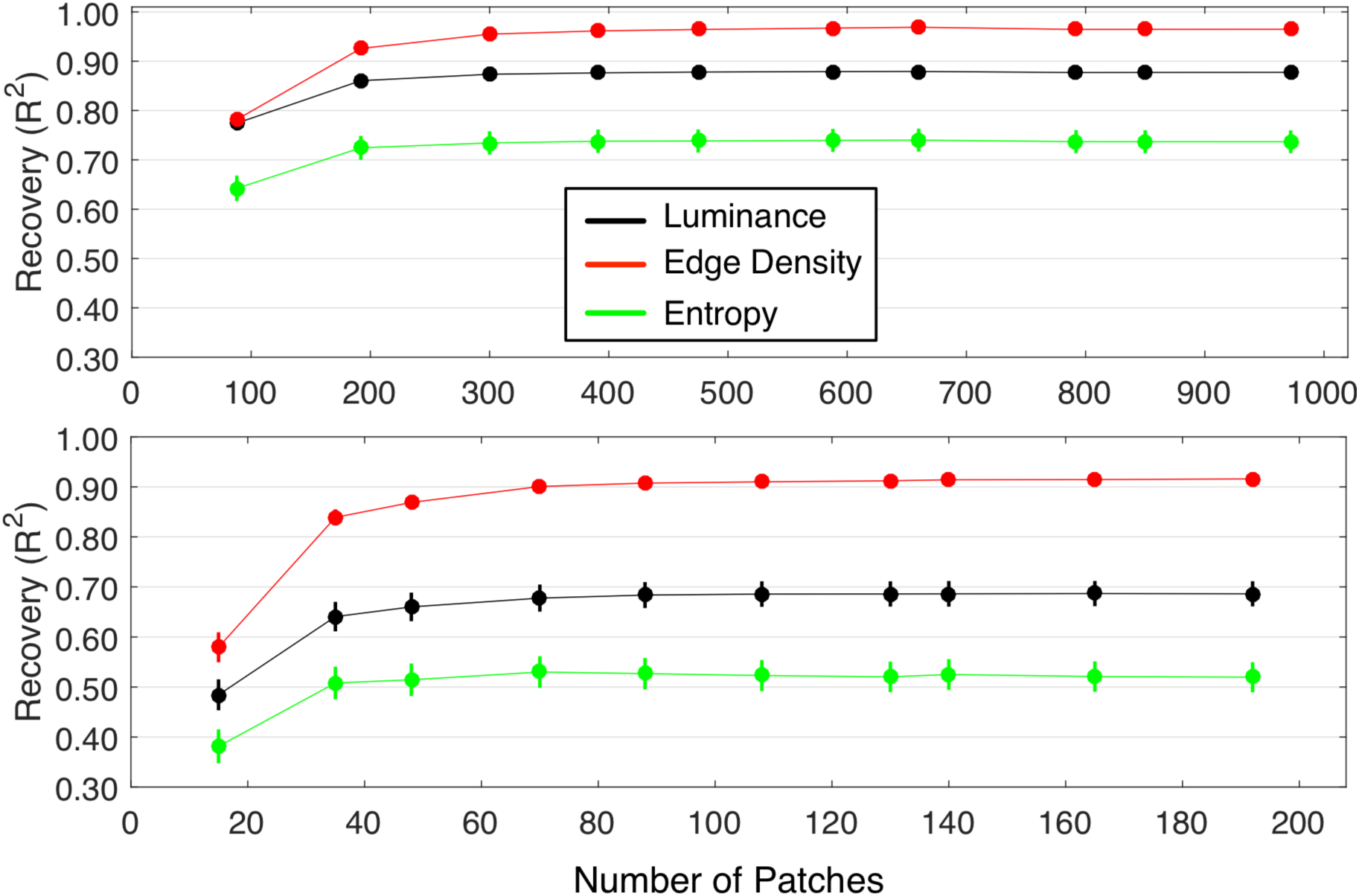
Ground truth recovery as a function of patch density for 3° and 7° patch sizes. The top panel shows the ground truth recovery (R2) across all 40 scenes for luminance, edge density, and entropy for the 3° patch size. The bottom panel shows the corresponding ground truth recovery (R^2^) for the 7° patch size. Error bars represent standard error of the mean.

It is worth noting that the recovery procedure makes two assumptions. First, it assumes that meaning can be interpolated from sub-sampling similarly to luminance, edge density, and entropy. Second, it assumes that our rating task provides an accurate estimate of meaning at each patch sample location. A priori we did not know whether these assumptions about meaning or our rating task were satisfied. While we still do not know whether the selected patch densities or rating task are optimal for measuring meaning, the accuracy of the meaning map prediction results suggests that the recovery simulations using image features provided reasonable sample density values for each patch size, and that the rating task provided reasonably accurate estimates of patch meaning.

